# Sex-specific sequels of early life stress on serine/threonine kinase activity in visceral adipose tissue from obese mice

**DOI:** 10.1101/2024.04.03.587852

**Authors:** Jacqueline Leachman, Justin Creeden, Meghan Turner, Nermin Ahmed, Carolina Dalmasso, Analia S. Loria

**Affiliations:** Department of Pharmacology and Nutritional Sciences, University of Kentucky, Lexington KY 405362; The Department of Neurosciences at the University of Toledo Medical Center

**Keywords:** ACE, BMI, Kinome, sex differences

## Abstract

Adverse childhood experiences (ACEs) are an established independent risk factor for chronic disease including obesity and hypertension; however, only women exposed to multiple ACEs show a positive relationship with BMI. Our lab has reported that maternal separation and early weaning (MSEW), a mouse model of early life stress, induces sex-specific mechanisms underlying greater blood pressure response to a chronic high fat diet (HF). Specifically, female MSEW mice fed a HF display exacerbated perigonadal white adipose tissue (pgWAT) expansion and a metabolic syndrome-like phenotype compared to control counterparts, whereas hypertension is caused by sympathoactivation in male MSEW mice. Thus, this study aimed to determine whether there is a sex-specific serine/threonine kinase (STKA) activity in pgWAT adipose tissue associated with early life stress. Frozen pgWAT was collected from MSEW and control, male and female mice fed a HF to assess STKA activity using the Pamstation12 instrument. Overall, MSEW induces significant reduction of 7 phosphokinases (|Z| >=1.5) in females (QIK, MLK, PKCH, MST, STE7, PEK, FRAY) and 5 in males (AKT, SGK, P38, MARK, CDK), while 15 were downregulated in both sexes (DMPK, PKA, PKG, RSK, PLK, DYRK, NMO, CAMK1, JNK, PAKA, RAD53, ERK, PAKB, PKD, PIM, AMPK). This data provides new insights into the sex-specific dysregulation of the molecular network controlling cellular phosphorylation signals in visceral adipose tissue and identifies possible target phosphokinases implicated in adipocyte hypertrophy as a result of exposure to early life stress. Identifying functional metabolic signatures is critical to elucidate the underlying molecular mechanisms behind the sex-specific obesity risk associated with early life stress.

## Introduction

The obesity epidemic in the United States of America affects 42.4% of the population regardless of sex; however, age adjustments show that severe obesity is 9.2% higher in women [1]. Additionally, the current statistics project that 50% of adults will have obesity by 2030 [2]. Adults with obesity typically have multiple organ system complications and, as a result, are more at risk for heart disease, stroke, type 2 diabetes, and multiple types of cancers [3-6]. An important limiting factor in controlling this epidemic is related to the unsuccessful long-term results in the clinical management of obesity [7, 8]. As there is a need for further investigation into the potential causes that may dampen the efficacy of treatments to prevent or reduce obesity, identifying modifiable risk factors affecting men and women’s obesity trajectories becomes a key step to addressing health disparities.

The development of obesity is a complex process influenced by genetics and environmental factors [9-11]. Lifestyle, nutrition, psychosocial environment, and socioeconomic status have been shown to shape the well-being of adult individuals, showing extremely detrimental consequences when adverse experiences occur during the first decade of life [12-14]. Notably, childhood obesity and the psychosocial influence of racial and ethnic disparities at a young age may have an unforeseen contribution to the obesity epidemic [15, 16]. Current growth trajectories predict that over half of toddlers and children will be obese by the age of 35 [17]. Supporting these trajectories, clinical and experimental studies have shown that prenatal life and early childhood determine the predisposition of the individual to gain weight and/or develop impairments in energy metabolism homeostasis [18, 19]. Therefore, any early life event that prevents the development of positive nutritional and lifestyle habits during childhood, including the influence of caregivers modeling healthy behaviors, may have a life-long deleterious impact on weight trajectories [20-22].

Protein kinases (a total 518 known in the human kinome) are the key enzymes catalyzing phosphorylation, a posttranslational modification that participates in the regulation of biological processes and molecular functions, such as insulin signal transduction and glucose disposal [23]. PamGene has developed a unique technology to perform real-time measurement of kinase activity.

We have shown previously that maternal separation and early weaning (MSEW) a model of neglect, combined with an unhealthy diet high in fat content mirrors the increased risk of developing obesity-induced hypertension [24-27]. Thus, this study aims to investigate the effects of early life stress on the kinome activity of adipose tissue from male and female MSEW mice fed a high-fat diet.

## Methods

### Animal model

Maternal Separation and Early Weaning (MSEW) was performed using C57BL/6J breeders given ad libitum access to food (2918 Teklad irradiated Global 18% Protein Rodent Diet) and water (Lexington city tap water treated by reverse osmosis). Animal rooms were maintained at 21±2 ºC and kept on a 14:10 light:dark cycle. MSEW litter pups were separated from the dam for 4 hours a day from postnatal days (PD) 2 to 5 and for 8 hours a day from PD 6 to 16 (21, 22, 64, 65). Pups were weaned early on PD 17. Separation occurred at the same time of day, during which the mother was removed from the cage and pups were placed into a clean cage in an incubator (30±1 ºC, 60% humidity). Normally reared litters served as Controls and remained undisturbed with the dam until weaning at PD 21.

### Animal Diet

Male and Female mice were placed onto a high fat diet (HF; D12492, 60% Kcal from fat, Research Diets) for 20 weeks.

### Experimental design

At week 20, mice were euthanized via exsanguination under anesthesia (ketamine/xylazine 100/10 mg/kg ip) to obtain pgWAT (epididymal fat in males and periovaric fat in females) for analysis of STKA. Samples from Control male, MSEW male, control female, MSEW female were pooled (n=8/group) and run in triplicate. Frozen tissue (50 mg) was homogenized to a final concentration of 25 ug/25ul.

### Kinase Activity

The Pamstation12 instrument (PamGene International; ‘s-Hertogenbosch, The Netherlands) was used to provide profiling of kinase activity in pgWAT samples. The device is loaded with serine/threonine microarray chips. Each chip has 4 wells that can be loaded with samples on each chip, and the Pamstation12 can accommodate 3 chips per run as shown in **Figure 1A**. 3 technical replicates of the 4 groups were loaded onto each chip to limit between-chip variation. The microarray used 144 (STK chip) reporter peptides from known serine/threonine kinase phosphorylation sites and measured the degree of phosphorylation in real-time by detecting fluorescently labeled antibodies at different exposure times.

**Figure 1:**
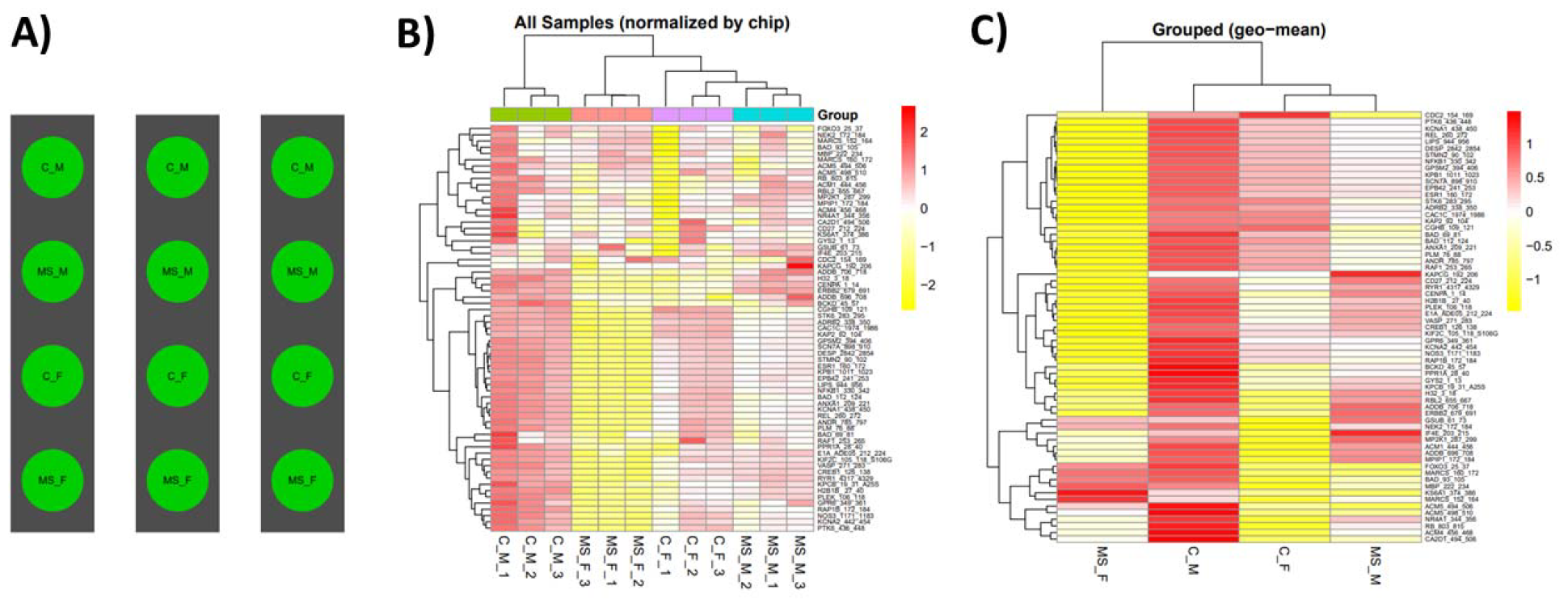
PamGene STK array analysis. Sample allocation is visualized across three PamGene STK chips (A). Signal intensities are of differentially phosphorylated peptides for each sample are visualized in a heatmap (B). Signal intensities averaged from technical replicates visualized in a heatmap (C).

### Image Analysis

Images taken by the PamStation of each array at different exposure times were converted to numerical values using Bionavigator (Pamgene). The software recognizes array grids with the aid of the searching algorithm (Pamgrid) to correctly identify each spot on the array. The numbers produced by this software represent the median value of the foreground pixels minus the median value of the background pixels to produce the median signal minus background.

### Data Tidying and Modeling

Raw data was transformed for analysis, modeling, and visualization by combining values from different exposure times, a simple linear regression model of the Medain_SigmBg as a function of exposure time is fitted. The slope of the model fit and R2 was used for quality control and samples comparison. The slope was also scaled by multiplying by 100 and log2 transformed.

### Global Signal Intensity

For a global signal intensity across all samples/groups, a heatmap was constructed based on the Slope_Transformed values. The heatmap represents all peptides present on the chip except for the positive/internal controls (pVASP_150_164, pTY3H_64_78, ART_025_CXGLRRWSLGGLRRWSL). The heatmap was scaled by row to highlight the peptides signals differences across the samples. A hierarchical unsupervised clustering was applied to peptides and samples to group similar signatures.

### Group Comparison

To compare between samples (MS_M vs C_M and MS_F vs C_F, a two-group comparison was performed. The Slope_Transformed ratio between each group, paired by chip, was calculated to represent the fold change. Only the peptides that pass the fold change threshold are considered significant. Also, quality control steps applied in each comparison to filter for peptides that do not hold to the criteria:

1. The Medain_SigmBg at max exposure 200ms must be equal or above 5
2. R2 of the linear model fit must be above or equal 0.9
3. Log fold change (LFC) cutoffs at (0.2, 0.3, 0.4)

## Results

### Data Acquisition and Transformation

To identify kinase pathways that may contribute to the sex-specific pathogenesis of early life stress in mice, we quantified the activity of 144 serine/threonine kinase in control and MSEW male and female mice. Figure 1A presents the run design for the three microarray chips used in this experiment. This schematic visually depicts the four groups loaded per chip, with three technical replicates to reduce variability. Figure 1B shows a heatmap of differentially phosphorylated peptides, normalized by chip and organized by unsupervised hierarchal clusters. The geometric means of the global signal intensities for each group are then clustered again in Figure 1C.

### Male Kinase Activity

Filtering parameters were applied to the male data, revealing 49 peptide “hits”, which were carried forward in the analysis. Hypophosphorylation was generally observed across pgWAT samples of MS males compared to control males, as indicated on the heatmap (Figure 2A). The signal intensities of these 49 hits are compared between control and MS males using a linear-fit model (Figure 2B). Figure 2C shows the waterfall plot for these peptides across each of the three technical replicates. Upstream analysis predicts kinases that can phosphorylate each peptide hit. These kinases are then plotted according to the log fold change of their corresponding peptides in a reverse KRSA plot (Figure 2D). MS male mice had decreased predicted activity of 13 kinases (Z>=2): protein kinase B (AKT, also known as PKB), AMP-activated protein kinase (AMPK), calcium/calmodulin-dependent protein kinase type 1 (CAMK1), dystrophia myotonica protein kinase (DMPK), Dual specificity tyrosine-phosphorylation-regulated kinase (DYRK), Extracellular signal-regulated kinase (ERK), c-Jun N-terminal kinases (JNK), protein kinase A (PKA), cGMP-dependent protein kinase or protein kinase G (PKG), Polo-like kinase (PLK), checkpoint kinase 2 (RAD53, also known as CHK2), ribosomal s6 kinase (RSK) and serum/glucocorticoid-regulated kinase (SGK).

**Figure 2:**
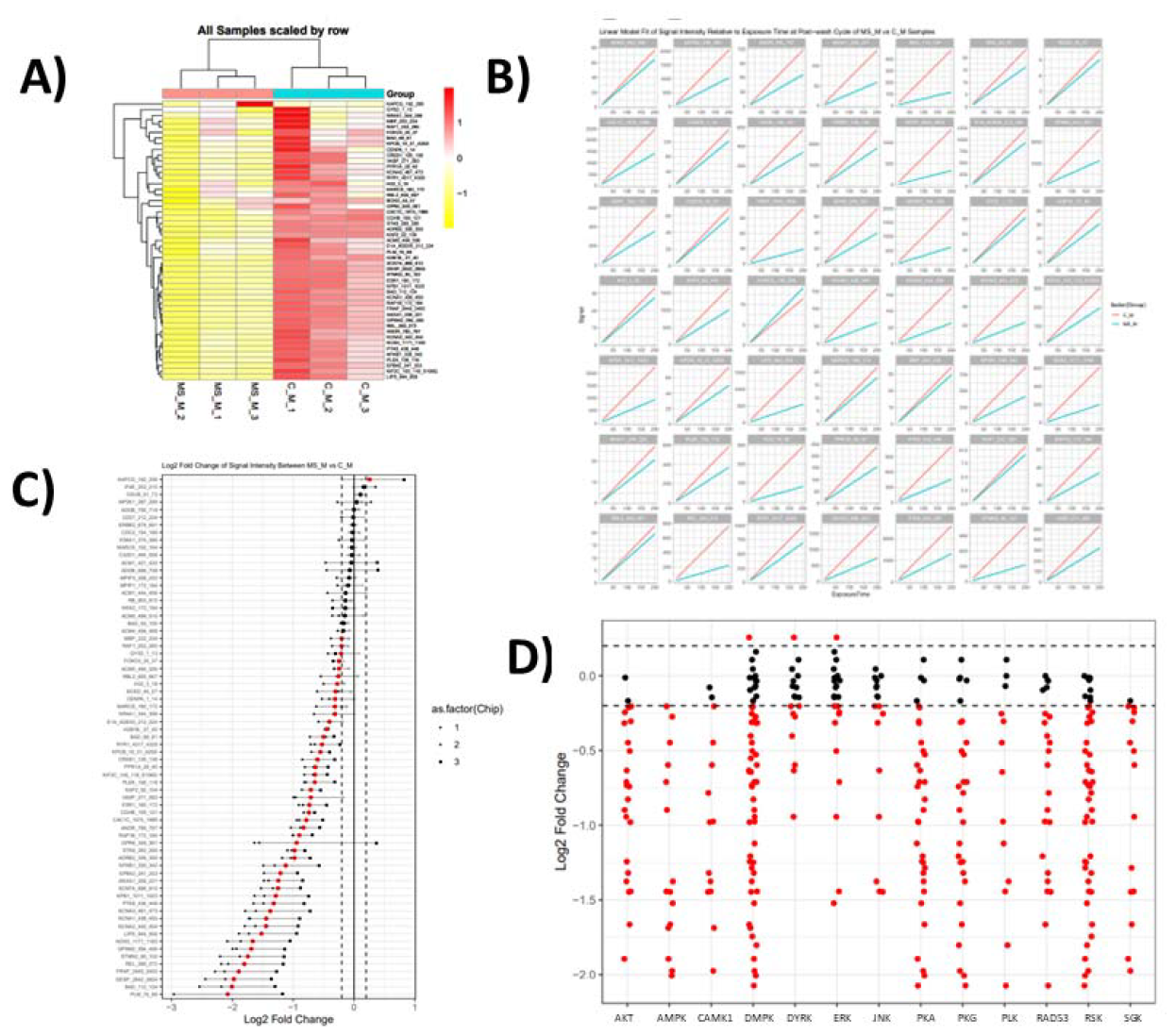
Analysis and prediction of differentially active kinases in males. Heat map of peptide phosphorylation signal intensities from MSEW and C male adipose tissue samples (A). MSEW and C signal intensities for 49 peptide hits displayed using linear-fit model (B). Waterfall plot of peptide hits showing variation across 3 technical replicates (C). Reverse KRSA plot of predicted differentially-active kinases in MSEW male adipose tissue (D).

### Female Kinase Activity

Filtering parameters were applied separately to female data, revealing 50 peptide hits for further analysis. The heatmap reveals both hypo-and hyperphosphorylated peptides (Figure 3A). A comparison of signal intensities for these 50 hits are between control and MS females is shown in Figure 3B using a linear-fit model. A waterfall plot of these hits are shown in Figure 3C. Upstream kinase analysis predicted 11 potential kinases, which are plotted in Figure 3D on a reverse KRSA plot. MS female mice had decreased predicted activity of the following 11 kinases (Z>=2): Calcium/calmodulin-dependent protein kinase type 1 (CAMK1), DMPK, DYRK, ERK, JNK, p21-activated kinase 1 (PAKA, also known as PAK1), PKA, PKG, PLK, RAD53, and RSK.

**Figure 3:**
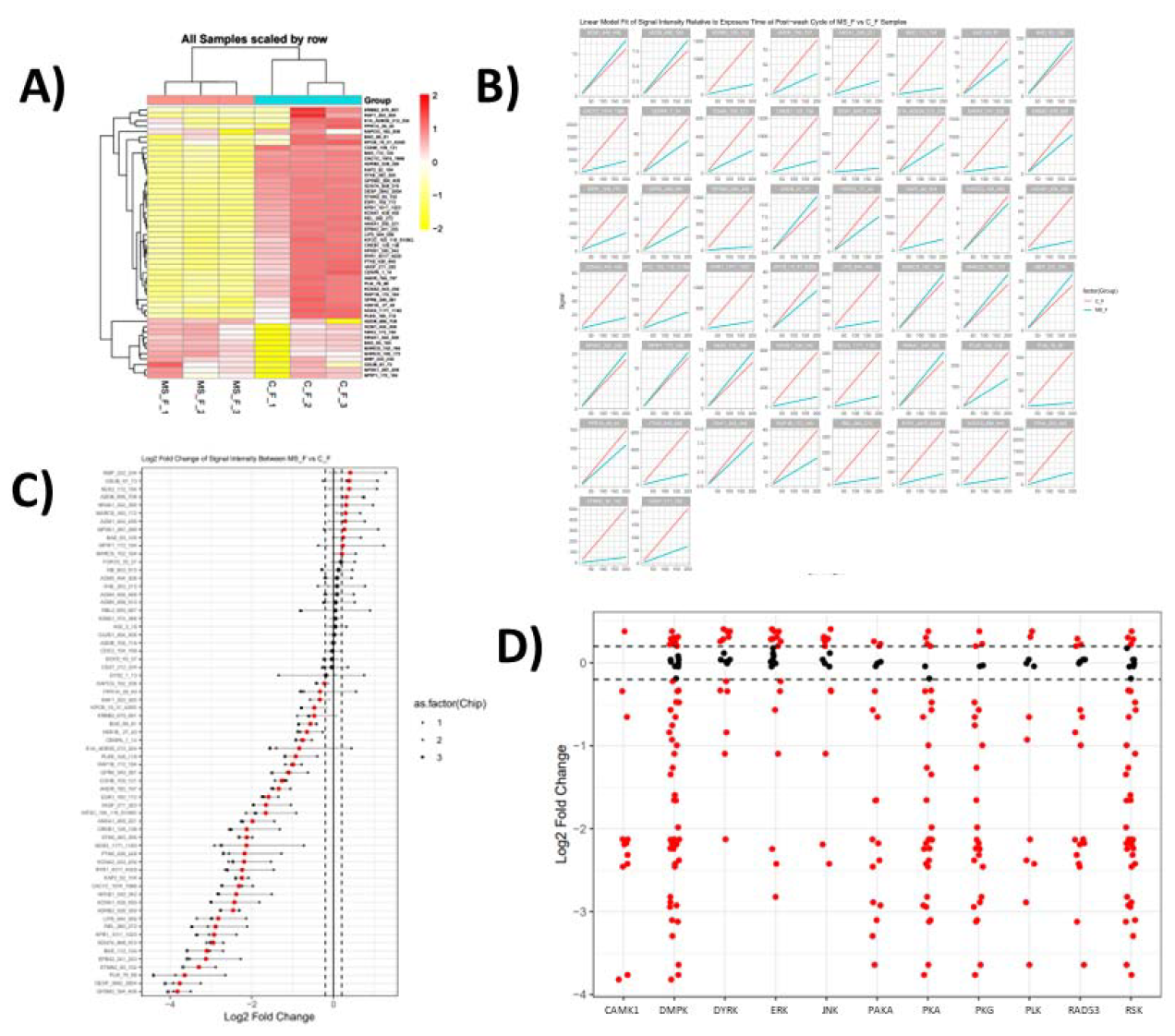
Analysis and prediction of differentially active kinases in females. Heat map of peptide phosphorylation signal intensities from MSEW and C female adipose tissue samples (A). MSEW and C signal intensities for 51 peptide hits displayed using linear-fit model (B). Waterfall plot of peptide hits showing variation across 3 technical replicates (C). Reverse KRSA plot of predicted differentially-active kinases in MSEW female adipose tissue (D).

### Upstream Kinase Analysis

Figure 4 visually depicts kinases that are predicted to be differentially activated in the pgWAT of in male MS, female MS, or ambiguously in MS mice. With the most stringent Z core threshold of 2 (Figure 4A), male MS mice have significantly different activation of AKT, AMPK, and SGK, while females differ only in PAKA activity. The MS protocol differentially activates DMPK, DYRK, PKA, ERK, RSK, JNK, RAD53, NMO, PKG, CAMK1, and PLK in the pgWAT of both sexes. At the threshold of Z>=1.75 (Figure 4B), males additionally show a difference in the activation of P38 and PIM. PAKA becomes ambiguously significant, along with PAKB, while female pgWAT shows differential activation of QIK, MLK, PKCH, MST, STE7, and PKD. With the lowest analytical threshold (Z>=1.5; Figure 4C), AMPK and PKD become significant between both sexes, and are joined by PIM. Male mouse pgWAT shows the additional differential activation of MARK and CDK, and female mouse pgWAT shows differential activation of PEK and FRAY in addition to the more stringent analysis.

**Figure 4:**
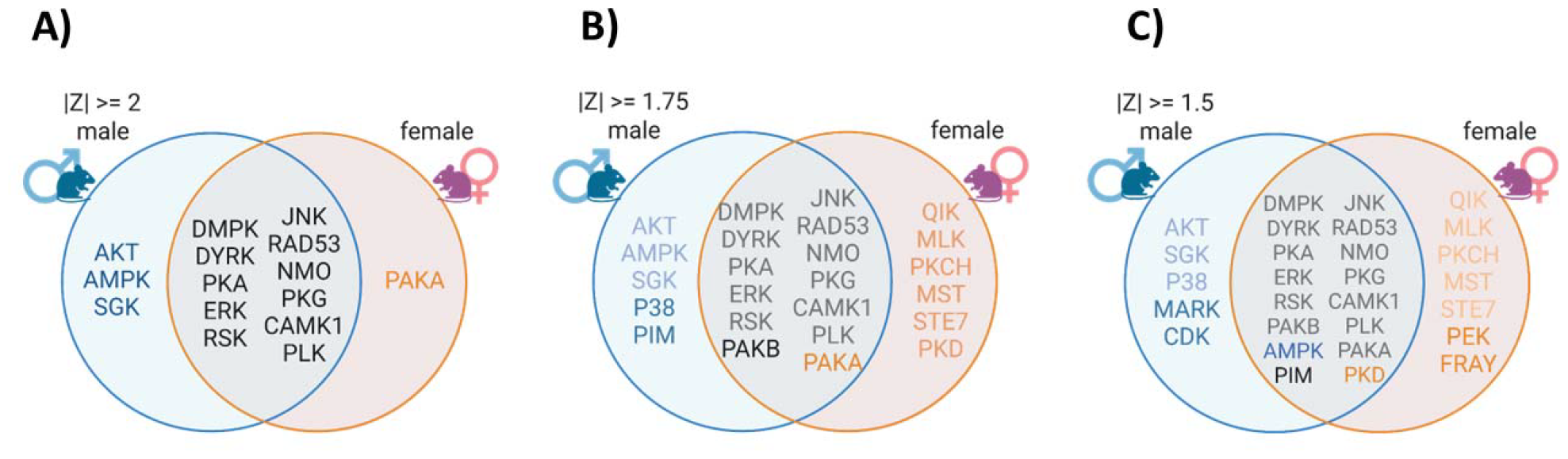
Differentially-activated kinases in male and female MSEW and C adipose tissue samples. Venn diagram of kinases predicted by KRSA analysis to be differentially-activated in male and female mouse adipose tissue at a Z-score significance of 2 (A), 1.75 (B), and 1.5 (C).

## Remarks

Our study is the first to show that early life stress induces sex-specific downregulation of STKA in adipose tissue from obese mice exposed to early life stress. We report that there are specific STKA signatures in male and female mice. Notably, female MSEW mice display fat expansion and increased adipokines in response to an obesogenic diet, while male MSEW mice show similar adiposity and circulating leptin levels compared to controls.

Calcium is a known significant secondary messenger in the process of adipogenesis by increasing the transcriptional activity of PPARy and C/EBPa [28]. DMPK is the most significant differentially active predicted kinase in our analysis and is implicated in calcium homeostasis by regulating both extracellular and intracellular calcium flux. For example, phosphorylation of subunit b of L-type calcium channels in skeletal muscle increases the voltage-sensitivity to extracellular calcium influx [29]. DMPK^-/-^ myotubules have smaller and slower calcium transients in response to acetylcholine stimulation compared to wild type, suggesting that phosphorylation by DMPK is necessary for adequate calcium signaling [29]. Phosphorylation of phospholamban in cardiomyocytes by DMPK facilitates uptake of Ca^2+^ to the sarcoplasmic reticulum, restoring intracellular calcium levels to steady-state [30]. Although studies have elucidated clear cellular mechanisms in calcium homeostasis for this kinase in myocytes, it’s specific role in adipocytes is currently unexplored. L-type calcium channels in rat adipocytes have, however, been linked to both steady-state and catecholamine-induced lipolysis [31, 32]. CAMK1 family kinases have diverse actions in response to calcium influx, and are differentially activated in our models as well[33].

The mitogen-activated protein kinase (MAPK) family of kinases is involved in diverse responses to physiological stressors such as DNA damage, oxidative stress, and infection. All three mammalian MAPK sub-families; JNK, P38, and ERK; are differentially activated by the MS protocol in adipose tissue of HFD-fed mice. JNKs have known involvement in obesity-induced insulin resistance [34], and downstream activation of RSK is a negative regulator of the insulin receptor, leading to insulin resistance[35]. ERK signaling promotes PPARy and C/EBPa transcriptional activity and is necessary for the adipogenic process [36]. Additionally, DYRK family kinases are known to be regulators of adipogenesis [37].

Additional biochemical validation is required to elucidate the contribution of altered kinases activity to the cardiometabolic phenotype displayed by male and female mice exposed to early life stress.

## References

1. Hales, C.M., et al., Prevalence of Obesity and Severe Obesity Among Adults: United States, 2017-2018. NCHS Data Brief, 2020(360): p. 1–8.

2. Ward, Z.J., et al., Projected U.S. State-Level Prevalence of Adult Obesity and Severe Obesity. N Engl J Med, 2019. 381(25): p. 2440–2450.

3. Kotsis, V., et al., Obesity and cardiovascular risk: a call for action from the European Society of Hypertension Working Group of Obesity, Diabetes and the High-risk Patient and European Association for the Study of Obesity: part A: mechanisms of obesity induced hypertension, diabetes and dyslipidemia and practice guidelines for treatment. J Hypertens, 2018. 36(7): p. 1427–1440.

4. Van Gaal, L.F., I.L. Mertens, and C.E. De Block, Mechanisms linking obesity with cardiovascular disease. Nature, 2006. 444(7121): p. 875–80.

5. Kahn, S.E., R.L. Hull, and K.M. Utzschneider, Mechanisms linking obesity to insulin resistance and type 2 diabetes. Nature, 2006. 444(7121): p. 840–6.

6. Iyengar, N.M., et al., Obesity and Cancer Mechanisms: Tumor Microenvironment and Inflammation. J Clin Oncol, 2016. 34(35): p. 4270–4276.

7. Montesi, L., et al., Long-term weight loss maintenance for obesity: a multidisciplinary approach. Diabetes Metab Syndr Obes, 2016. 9: p. 37–46.

8. Hall, K.D. and S. Kahan, Maintenance of Lost Weight and Long-Term Management of Obesity. Med Clin North Am, 2018. 102(1): p. 183–197.

9. Maes, H.H., L.J. Neale Mc Fau - Eaves, and L.J. Eaves, Genetic and environmental factors in relative body weight and human adiposity. 1997(0001-8244 (Print)).

10. Hinney, A.A.-O., A.A.-O. Körner, and P.A.-O. Fischer-Posovszky, The promise of new anti-obesity therapies arising from knowledge of genetic obesity traits. 2022(1759-5037 (Electronic)).

11. Locke, A.E., et al., Genetic studies of body mass index yield new insights for obesity biology. 2015(1476-4687 (Electronic)).

12. Anda, R.F., et al., Adverse childhood experiences and chronic obstructive pulmonary disease in adults. 2008(0749-3797 (Print)).

13. Power, C. and C. Hertzman, Social and biological pathways linking early life and adult disease. 1997(0007-1420 (Print)).

14. Shonkoff, J.P., B.S. Boyce Wt Fau - McEwen, and B.S. McEwen, Neuroscience, molecular biology, and the childhood roots of health disparities: building a new framework for health promotion and disease prevention. 2009(1538-3598 (Electronic)).

15. Pan, L., et al., Trends in Obesity Among Participants Aged 2-4 Years in the Special Supplemental Nutrition Program for Women, Infants, and Children - United States, 2000-2014. MMWR Morb Mortal Wkly Rep, 2016. 65(45): p. 1256–1260.

16. Petersen, R., L. Pan, and H.M. Blanck, Racial and Ethnic Disparities in Adult Obesity in the United States: CDC’s Tracking to Inform State and Local Action. Prev Chronic Dis, 2019. 16: p. E46.

17. Ward, Z.J., et al., Simulation of Growth Trajectories of Childhood Obesity into Adulthood. N Engl J Med, 2017. 377(22): p. 2145–2153.

18. Dietz, W.H., Critical periods in childhood for the development of obesity. Am J Clin Nutr, 1994. 59(5): p. 955–9.

19. Danese, A. and M. Tan, Childhood maltreatment and obesity: systematic review and meta-analysis. Molecular Psychiatry, 2014. 19(5): p. 544–554.

20. Birch, L., J.S. Savage, and A. Ventura, Influences on the Development of Children’s Eating Behaviours: From Infancy to Adolescence. Can J Diet Pract Res, 2007. 68(1): p. s1–s56.

21. Lytle, L.A., et al., How do children’s eating patterns and food choices change over time? Results from a cohort study. Am J Health Promot, 2000. 14(4): p. 222–8.

22. Mannino, M.L., et al., The quality of girls’ diets declines and tracks across middle childhood. Int J Behav Nutr Phys Act, 2004. 1(1): p. 5.

23. Chen, J., et al., Protein kinases in cardiovascular diseases. 2022(2542-5641 (Electronic)).

24. Dalmasso, C., et al., Sensory signals mediating high blood pressure via sympathetic activation: role of adipose afferent reflex. American Journal of Physiology-Regulatory, Integrative and Comparative Physiology, 2020. 318(2): p. R379–R389.

25. Murphy, M.O., et al., A model of neglect during postnatal life heightens obesity-induced hypertension and is linked to a greater metabolic compromise in female mice. International Journal of Obesity, 2018. 42(7): p. 1354–1365.

26. Leachman, J.R., et al., Exacerbated obesogenic response in female mice exposed to early life stress is linked to fat depot-specific upregulation of leptin protein expression. Am J Physiol Endocrinol Metab, 2020. 319(5): p. E852–E862.

27. Leachman, J.R., et al., Early life stress exacerbates obesity in adult female mice via mineralocorticoid receptor-dependent increases in adipocyte triglyceride and glycerol content. Life Sci, 2022. 304: p. 120718.

28. Zhai, M., et al., Involvement of calcium channels in the regulation of adipogenesis. Adipocyte, 2020. 9(1): p. 132–141.

29. Benders, A.A., et al., Myotonic dystrophy protein kinase is involved in the modulation of the Ca2+ homeostasis in skeletal muscle cells. Journal of Clinical Investigation, 1997. 100(6): p. 1440–1447.

30. Kaliman, P., et al., Myotonic Dystrophy Protein Kinase Phosphorylates Phospholamban and Regulates Calcium Uptake in Cardiomyocyte Sarcoplasmic Reticulum. Journal of Biological Chemistry, 2005. 280(9): p. 8016–8021.

31. Zhang, F., et al., Calcium Supplementation Enhanced Adipogenesis and Improved Glucose Homeostasis Through Activation of Camkii and PI3K/Akt Signaling Pathway in Porcine Bone Marrow Mesenchymal Stem Cells (pBMSCs) and Mice Fed High Fat Diet (HFD). Cellular Physiology and Biochemistry, 2018. 51(1): p. 154–172.

32. Fedorenko, O.A., et al., CaV1.2 and CaV1.3 voltage-gated L-type Ca2+ channels in rat white fat adipocytes. Journal of Endocrinology, 2020. 244(2): p. 369–381.

33. Colomer, J. and A.R. Means, Physiological roles of the Ca2+/CaM-dependent protein kinase cascade in health and disease. Subcell Biochem, 2007. 45: p. 169–214.

34. Yung, J.H.M. and A. Giacca, Role of c-Jun N-terminal Kinase (JNK) in Obesity and Type 2 Diabetes. Cells, 2020. 9(3).

35. Smadja-Lamère, N., et al., Insulin activates RSK (p90 ribosomal S6 kinase) to trigger a new negative feedback loop that regulates insulin signaling for glucose metabolism. J Biol Chem, 2013. 288(43): p. 31165–76.

36. Prusty, D., et al., Activation of MEK/ERK Signaling Promotes Adipogenesis by Enhancing Peroxisome Proliferator-activated Receptor γ (PPARγ) and C/EBPα Gene Expression during the Differentiation of 3T3-L1 Preadipocytes. Journal of Biological Chemistry, 2002. 277(48): p. 46226–46232.

37. Masaki, S., et al., Design and synthesis of a potent inhibitor of class 1 DYRK kinases as a suppressor of adipogenesis. Bioorg Med Chem, 2015. 23(15): p. 4434–4441.

